# *Bacillus subtilis* maintains antibiofilm activity against *Staphylococcus aureus* following adaptive laboratory evolution

**DOI:** 10.1101/2023.10.15.562438

**Authors:** Kyle R. Leistikow, Elisabeth Solis, Jehan Khaled, Christopher W. Marshall, Krassimira R. Hristova

## Abstract

Quorum sensing interference has been touted as an ideal mechanism for the development of new anti-virulence therapies. Recent work has established *Bacillus subtilis* 6D1, a Gram-positive spore forming bacterium with probiotic qualities, produces metabolites that inhibit *Staphylococcus aureus* virulence and biofilm formation via quorum sensing interference. However, it remains unknown how long-term exposure to these molecules drive *S. aureus* adaptation and evolution. *S. aureus* planktonic cells and biofilms were propagated in the presence of *B. subtilis* 6D1 cell free extracts (CFE) for ∼73 generations. Fitness, virulence, and antibiotic resistance assays of the ancestor and all evolved lineages revealed the emergence of treatment and lifestyle associated ecological traits. Compared to the ancestor and media-evolved lineages, *S. aureus* lineages evolved in the presence of *B. subtilis* 6D1 CFE were less competitive in a biofilm and exhibited increased phenotypic sensitivity to multiple antibiotics. Notably, *B. subtilis* 6D1 CFE maintained its ability to inhibit *S. aureus* biofilm growth and disassemble mature biofilm in all evolved lineages. *S. aureus* populations propagated in the presence of CFE acquired missense mutations in genes associated with plasmid-borne efflux systems and RNA polymerase. Furthermore, CFE-evolved lineages did not develop mutations in both competence and drug resistance pathways found in similarly evolved control lineages. Our data suggest long-term exposure to biofilm inhibitory molecules, like those produced by *B. subtilis* 6D1, can reduce *S. aureus’* fitness in a biofilm and increase sensitivity to multiple antibiotics.

**Importance:** Quorum sensing interference (QSI) has been touted as an ideal mechanism to diminish bacterial virulence and improve antibiotic killing, however few studies investigate the genetic and phenotypic adaptations that occur after long-term exposure to QSI therapies. Recent studies revealed *Bacillus subtilis* reduces biofilm formation and virulence via signaling interference with the *S. aureus* Agr QS system; however, it remains unclear how long-term exposure to these compounds drives *S. aureus* adaptation and evolution. This study helps to address these gaps by investigating whether QSI strategies deployed by probiotic bacteria are viable approaches to increase antibiotic efficacy without increasing antibiotic resistance evolution.

## Introduction

Antibiotic resistant *Staphylococcus aureus* is a leading cause of pneumonia, sepsis, endocarditis, and soft tissue infections - made more difficult to treat due to its evolving resistance to front-line antibiotics(1, 2) and its ability to form biofilms(3, 4). Biofilms are an aggregated heterogenous community of cells surrounded by an extracellular polymeric matrix that can increase antibiotic resistance 1000-fold(5). Microbial competitive interactions that occur in a biofilm increase the likelihood of high frequency resistant cells through mutations or gene transfer(6), and in *S. aureus,* these competitive biofilm environments increase virulence(7, 8) and antibiotic resistance(9, 10). Additionally, microbial experimental evolution assays have shown microorganisms evolved in a biofilm develop unique phenotypes compared to planktonic cells(11, 12), including adaptations that increase virulence(13, 14) and alter the development of antibiotic resistance(15, 16). Therefore, if biofilms are disrupted or inhibited, virulence may also decrease(17, 18).

In *S. aureus*, biofilm formation and dispersal is controlled in part by the Agr quorum sensing (QS) system(19). It has been reported quorum sensing interference (QSI) can prevent the formation of biofilm structures and increase antibiotic killing(20, 21). Unlike traditional antibiotics, which mainly aim to inhibit and kill microorganisms, QSI agents do not typically alter the growth of microorganisms in planktonic conditions, nor are they believed to induce the development of antibiotic resistance since quorum-sensing genes are off-target for antibiotics(22, 23). Additionally, QS coordinates the production of many important extracellular factors that are cooperative “public goods” for the population. QS signal-blind mutants that do not produce these goods, but benefit from them, invade wild-type populations with negative frequency dependence - as their proportion increases in a population, their relative fitness decreases as there are fewer cooperators to exploit(24). This suggests QSI-resistant mutants are only fit when rare, so even if QSI-resistant mutants develop, they will likely not rise to fixation otherwise the population might collapse(25, 26). Given the therapeutic potential and presumed long-term feasibility of these compounds, it is essential that QSI strategies be studied in an evolutionary context, because even QSI can be overcome through adaptive evolution.

Resistance mechanisms against well-characterized quorum quenchers have been found in the laboratory as well as in clinical strains, demonstrating that resistance against these kinds of compounds is possible(27, 28). Fortunately, long-term experimental microbial evolution assays can provide unique opportunities to investigate how pathogenic populations adapt under long term selection of QSI compounds(29).

Probiotic bacterial species, namely *Bacillus subtilis,* produce an array of antimicrobials that can act synergistically to eliminate phylogenetically distinct competitors(30) and significantly slow down the selection of antibiotic resistance in Gram-negative pathogens(31). *B. subtilis* also produces QSI molecules capable of reducing *S. aureus* biofilm formation and host colonization(32–36), however it remains unknown how long-term exposure to these compounds drive *S. aureus* adaptation and evolution. Recent work in our lab has identified a single strain, *B. subtilis* 6D1, inhibits *S. aureus* biofilm formation and virulence and increases antibiotic sensitivity through an Agr-mediated QSI mechanism(37). The main objective of this work was to investigate how long-term exposure to these probiotic-derived compounds with QSI activity drive *S. aureus* evolution, fitness, virulence, and antibiotic resistance at the whole population level. Identifying the genes and the subsequent mutations that confer resistance to known QS pathways, like Agr, will provide a more comprehensive understanding of the modes of action of these new anti-virulence strategies and how resistance to them evolves.

## Methods

### Culture conditions and preparation of *B. subtilis* 6D1 cell free extracts

*B. subtilis* 6D1 cell free extracts were prepared as previously described(37). Briefly, exponential phase *B. subtilis* 6D1 cultures grown in trypticase soy broth (TSB) were individually cultured in 125mL baffled shake flasks containing 25mL TSB. Flasks were incubated at 37°C shaking at 230 RPM for 48 hours. After incubation, cultures were centrifuged at 4°C at 10,500 x g for 20 minutes. Cell free extract (CFE) was harvested and filter-sterilized using a 0.22μm Surfactant-free cellulose acetate (SCFA) filter (Corning, USA). CFEs were flash frozen in liquid nitrogen and stored at −80°C for future use.

### Adaptive laboratory evolution of *S. aureus* ATCC 29213

Daily passages of *S. aureus* ATCC 29213 planktonic or biofilm cultures were performed for ∼73 generations (11 days) resulting in a ∼99% probability of obtaining a mutation at any given site within the *S. aureus* ATCC 29213 genome(38, 39). This genome is comprised of a 2.76 Mb chromosome (GenBank sequence: CP078521.1) and 4.69 Kb plasmid (GenBank sequence: CP078522.1). Figure 1 outlines the experimental design. Briefly, five independently propagated *S. aureus* populations exposed to either 10% v/v PBS or 10% v/v *B. subtilis* 6D1 CFE were passaged daily into 5 mL of fresh 1.5% TSB + 0.3% glucose media(40, 41). Planktonic cultures were diluted 1:100 during each passage. In biofilm cultures, a single 7mm polystyrene bead was transferred daily to recapitulate the entire biofilm life cycle(11, 42). All cultures were incubated in a roller drum for 24 hours at 37°C between passages. Populations were preserved every other day and kept at −80°C for future use. Population sizes were determined via CFU plating on TSA plates after 1, 6, and 11 days.

**Figure 1:**
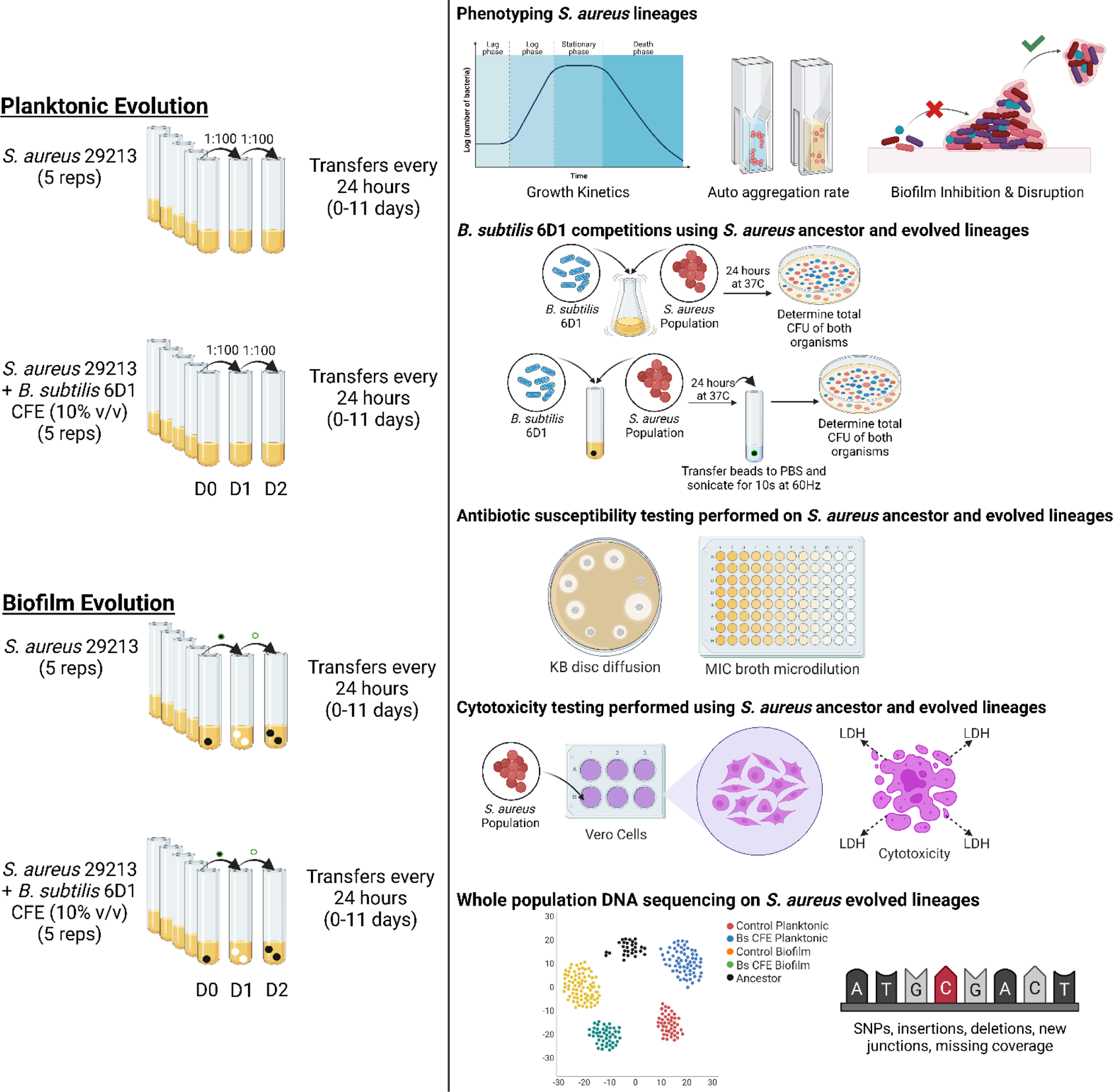
Experimental design investigating the ecological effects of *S. aureus* ATCC 29213 evolution after long-term exposure to *B. subtilis* 6D1 derived cell free extracts harboring QSI activity. CFE = Cell free extract; KB = Kirby-Bauer; MIC = Minimum inhibitory concentration; LDH = Lactate dehydrogenase; SNP = Single nucleotide polymorphism

### Growth assays

All lineages were revived 1:100 in evolution media (1.5% TSB + 0.3% glucose) overnight and then seeded at 5 x 10^5^ CFU in 200μL evolution media. Growth kinetics assays were performed in 96 well plates (Corning, USA) using a Multiskan GO spectrophotometer (Thermo Scientific, USA). Plates were incubated at 37°C in the instrument and orbitally shaken for 10 seconds prior to each read. Growth curves were subtracted from baseline based on each population’s initial turbidity. Endpoint cell density assays differed in that they were continuously shaken at 150 RPM in a 37°C incubator for 24 hours prior to reading turbidity. Each lineage was assayed in quadruplicate and averaged by treatment/lifestyle.

### Biofilm inhibition and disruption assays

*B. subtilis* 6D1 CFE (10% v/v) was applied to Nunc™ treated 96 well microplates (Thermo Scientific, USA) containing 1.5% TSB + 0.3% glucose. Test wells were seeded with 5 x 10^5^ CFU/mL of *S. aureus* and incubated statically at 37°C. After 24 hours, media was removed, and biofilms were gently washed three times with PBS, fixed for 20 minutes at 55°C, and stained with 0.1% crystal violet (CV) for 15 minutes. After 15 minutes, CV was removed, and wells were washed twice with PBS and allowed to dry. CV was solubilized in 30% acetic acid for 20 minutes. Biofilm inhibition was measured spectrophotometrically relative to untreated control wells at OD_595_. To assess biofilm disruption, 10% v/v *B. subtilis* 6D1 CFE was mixed with PBS and applied to *S. aureus* 24-hour mature biofilms and incubated at 37°C shaking at 100 RPM for 2 hours. To account for any mechanical disruption not driven by CFE, biofilms treated only with PBS were used as a negative control. Residual biofilms were fixed and quantified by crystal violet staining as described above. Each lineage was assayed in quintuplicate and averaged by treatment/lifestyle.

### Sedimentation assays

Sedimentation assays were conducted according to previous methods(43). To avoid the auto aggregative effects driven by extracellular exopolysaccharides, media composition, or pH(44, 45), all populations were standardized in Gibco™ PBS, pH 7.4 (Thermo Scientific, USA). Briefly, each evolved *S. aureus* population was revived in evolution media (1:100) and incubated at 37°C overnight at 200RPM. Each culture was then centrifuged for 20 minutes at 3200 x g, decanted, and standardized in PBS to ∼1 x 10^9^ CFU/mL (1.5 OD_600_). To ensure sedimentation rates were calculated comparably, each newly standardized PBS culture was centrifuged for an additional 20 minutes at 3200 x g, vortexed for 30 seconds and 1mL was pipetted into a cuvette. Turbidity (OD_600_) was measured immediately using a Multiskan GO spectrophotometer (Thermo Scientific, USA). Cuvettes were kept at room temperature between reads. Percent auto aggregation was expressed as a function of the culture turbidity at Time 0. Each lineage was assayed in triplicate in the presence and absence of 10% v/v *B. subtilis* 6D1 CFE and averaged per treatment/lifestyle.

### Competition assays

For planktonic competitions, 250μL (5 x 10^7^ CFU) of exponential phase *B. subtilis* 6D1 and *S. aureus* populations were added to 25mL TSB in a baffled shake flask and incubated at 37°C shaking at 200 RPM. Biofilm competitions utilized 7 mm polystyrene beads as a surface for cell attachment, biofilm growth, and biofilm dispersal(15). For biofilm competitions, equal concentrations of *B. subtilis* 6D1 and *S. aureus* populations (∼1 x 10^5^ – 1 x 10^6^) were added to a glass test tube containing 5mL 1.5% TSB + 0.3% glucose media(46) and a single polystyrene bead(11, 16). After 24 hours incubation at 37°C in a roller drum, the bead was placed in 1mL PBS and sonicated with a handheld Qsonica model CL-188 instrument (QSonica, USA) for 10 seconds at 60Hz to remove attached cells. For all competitions, CFU counts were obtained for both bacterial species at time point 0 and after 24 hours incubation. Strain fitness and selection rate were calculated as previously described by Travisano and Lenski(47).

### Cytotoxicity assays

CCL81 (Vero) cells were cultivated in Gibco™ Dulbecco’s Modified Eagle Medium (Thermo Scientific, USA) consisting of 10% fetal bovine serum and 0.1% Gibco™ Antibiotic-Antimycotic (Thermo Scientific, USA). Vero cells were cultured for 7 days at 37°C in a carbogen (95% O2, 5% CO2) atmosphere, with the culture medium being changed every other day until cells were confluent. Approximately 2 × 10^4^ cells were seeded into new 96 well plates (Corning, USA) and allowed to form monolayers for 48 hours prior to cytotoxicity testing. After removing media, cells were washed twice with Gibco™ PBS, pH 7.4 (Thermo Scientific, USA) and equilibrated in EC buffer (135mM NaCl, 15mM HEPES, 1mM MgCl_2_, 1mM CaCl_2_) for four hours before adding bacteria. Exponential phase *S. aureus* populations grown in 1.5% TSB + 0.3% glucose were standardized to OD_600_ 1.0, centrifuged, washed twice with PBS, and applied to test wells at a final concentration of ∼1.5 x 10^7^ CFU/well. Infected cell lines were incubated at 37°C in a carbogen environment for two hours prior to conducting cytotoxicity assays using a lactate dehydrogenase (LDH) kit (Abcam, UK) as previously described(48). After treatment, cells were centrifuged at 600g for 5 minutes. Cytotoxicity was normalized to 100% using a cell lysis positive control solution according to the manufacturer’s protocol. Three independent experiments were performed per *S. aureus* lineage and averaged by treatment/lifestyle. Cell lines were imaged at 20X magnification using a Nikon TMS inverted phase contrast microscope (Nikon Corporation, Japan).

### Antibiotic susceptibility testing

Both Kirby Bauer disc diffusion and broth microdilution assays were conducted on Mueller Hinton (MH) media (BD Life Sciences, USA) (49). To revive cultures, *S. aureus* populations were seeded in evolution media (1:100) and incubated at 37°C at 200 RPM. 1.5 x 10^8^ CFU/mL of exponential phase *S. aureus* population cultures (OD_600_= 0.6) were spread onto square MH agar plates and allowed to dry. Sensi-discs™ infused with known concentrations of nine different antibiotics (Table 1) were applied to the agar with sterile forceps and incubated at 37°C for 24 hours. The resulting zones of inhibition were measured (mm) and designated as susceptible, intermediate, or resistant according to the CLSI guidelines for *Staphylococcus*. Broth microdilutions were conducted using reduced antibiotic concentrations to assess differences in minimum inhibitory concentration (MIC) below those found in the infused discs. In these experiments, 100μL MH broth harboring different concentrations of antibiotics was inoculated with 5 x 10^5^ CFU/mL exponential phase *S. aureus* population cultures in 96-well polystyrene plates and incubated at 37°C at 150 RPM for 24 hours. All experiments were performed in triplicate and compared to the ancestor.

**Table 1.** Antibiotics and their mechanisms of action used to quantify *S. aureus* population sensitivity.

|  |  |  |  |
| --- | --- | --- | --- |
| **Antibiotic** | **Target** | **Class** | **Type** |
| Ceftazidime | Penicillin binding protein | Cephalosporin (3rd Generation) | Cell Wall Synthesis Inhibitors |
| Cefotaxime | Penicillin binding protein | Cephalosporin (3rd Generation) |  |
| Vancomycin | Peptidoglycan cross-linking | Glycopeptide |  |
| Gentamicin | Ribosome 30S subunit | Aminoglycoside | Protein Synthesis Inhibitors |
| Erythromycin | Ribosome 50S subunit | Macrolide |  |
| Tetracycline | Ribosome 30S subunit | Tetracycline |  |
| Sulfamethoxazole/Trimethoprim | Folate synthesis | Sulfonamide | Nucleic Acid Synthesis Inhibitors |
| Levofloxacin | DNA gyrase/topoisomerase IV | Fluoroquinolone |  |
| Rifampin | RNA polymerase | Ansamycin |  |

### Whole population genome sequencing

DNA was extracted from Day 11 populations (n=20, 5/treatment) using Qiagen DNeasy Blood and Tissue Kit (Qiagen, Germany) adapted from the manufacturer’s instructions to include a lysostaphin lysis step(50). Sample libraries were prepared using an Illumina DNA Prep kit and 10bp unique dual indices (Integrated DNA Technologies, USA). Whole population genome sequencing (650 Mbp) was performed on *S. aureus* populations using an Illumina NextSeq 2000. Sequence reads were demultiplexed and adapters were removed using bcl2fastq (2.20.0.445)(51). Variant calling analyses assessing differences in mutation presence and frequency between the *S. aureus* ancestor (GenBank Accession # ASM2287046v1) and evolved populations were performed on 2×151bp paired-end reads using breseq (v0.37.1) in polymorphism mode(52). These parameters call mutations only if they are present within the population at a frequency of at least 5% and are in at least two reads from each strand. Filtering, allele frequencies, and plotting were done in R software (v4.2.0; www.r-project.org) with the packages ggplot2 (v3.4.0; https://CRAN.R-project.org/package=ggplot2) and tidyr (https://CRAN.R-project.org/package=tidyr). Protein sequences harboring missense mutations were mapped onto Alphafold2(53) predicted structures and visualized using Pymol(54).

### Statistical analysis

Results are presented as mean ± standard error of the mean (SEM) unless noted otherwise. Statistical analyses were performed using GraphPad Prism 9.5.0 (San Diego, CA). Differences where p<0.05 were considered significant. Differences in biofilm inhibition were analyzed using a 2-tailed unpaired t-test. Sedimentation assays were analyzed using two-way ANOVA followed by Tukey’s multiple comparisons. Antibiotic susceptibility, Vero cell cytotoxicity, and competition results were analyzed using one-way ANOVA, followed by Tukey’s multiple comparisons. Differences in normalized plasmid read abundance were analyzed using a non-parametric Kruskal-Wallis test followed by Dunn’s multiple comparisons.

### Data availability

All sequencing reads were deposited in the NCBI sequence read archive under BioProject accession number PRJNA1006038. Detailed methods regarding data processing can be found at https://github.com/sirmicrobe/lab_breseq_workflow

## Results

### B. subtilis 6D1 CFE maintains antibiofilm activity against all evolved S. aureus populations regardless of treatment or lifestyle

Previous work has shown that *B. subtilis* 6D1 CFE inhibits biofilm growth of *S. aureus*(37), therefore, we wanted to test if *S. aureus* could evolve resistance to this antibiofilm activity and by what mechanism. To first assess how our evolutionary conditions affected population size and growth rate, population size was determined for all treatment replicates and averaged by treatment/lifestyle after 24 hours, and at the end of the 11-day directed evolution experiment. Neither the control nor the *B. subtilis* 6D1 CFE (Bs CFE)-exposed planktonic populations differed in population size throughout the experiment (Fig. 2A). However, Bs CFE-treated biofilm lineages significantly reduced population sizes from 7.19 ± 0.11 to 6.52 ± 0.13 log_10_ CFU/bead after 11 days (Fig. 2B). To avoid bottlenecking in biofilm populations as a result of CFE exposure, beads were not rinsed prior to population transfers or enumerations. Therefore, the high cell density observed on beads after CFE exposure can likely be attributed to an increased number of unattached cells being included in our CFU counts (Fig. S4). Despite these changes in population size, growth rate assays conducted in the same evolution media revealed mean doubling time decreased for all lineages compared to the ancestor, an indication that evolved populations adapted to the growth media (Table S1).

**Figure 2.**
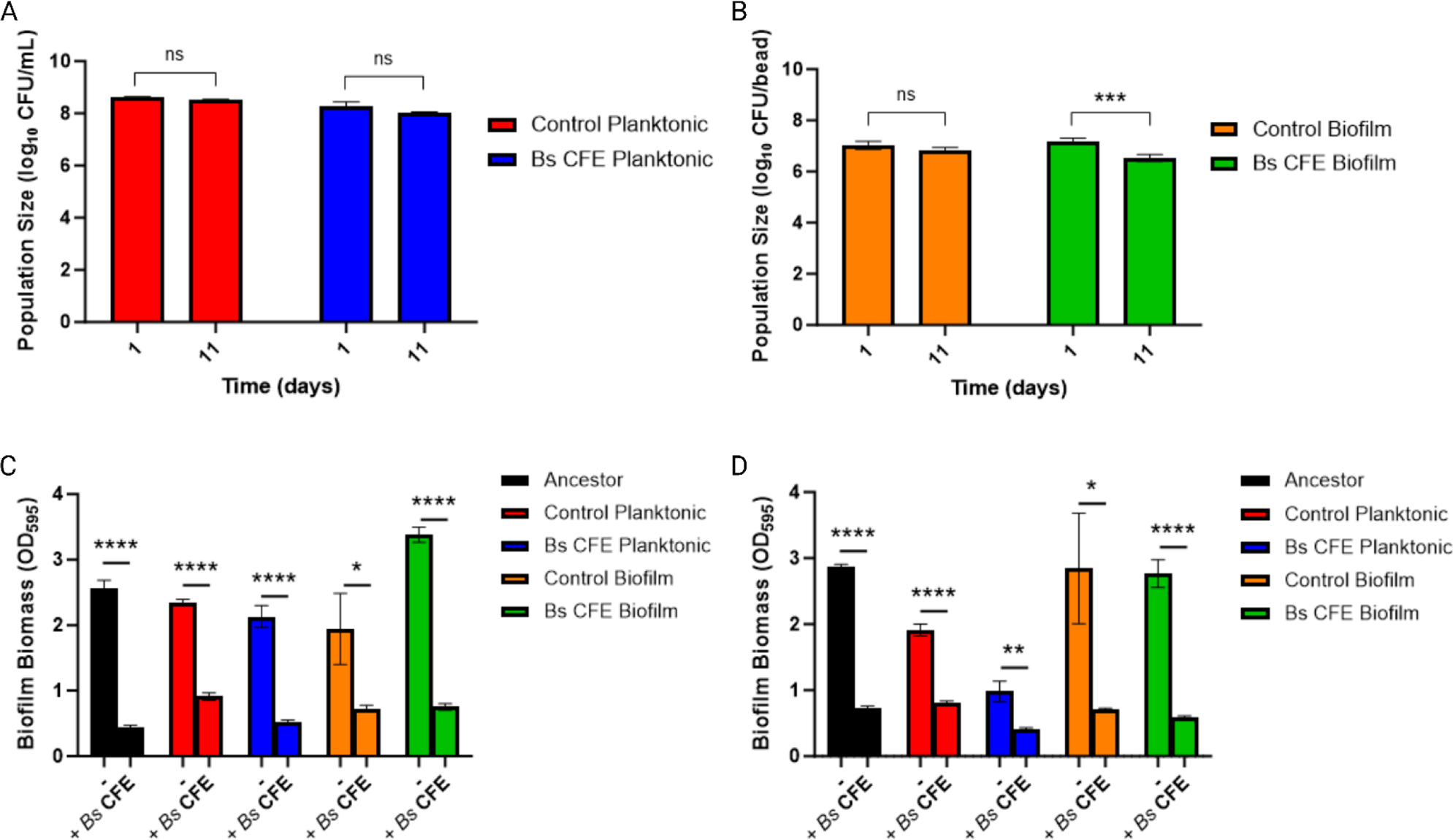
*B. subtilis* 6D1 CFE maintains antibiofilm activity against all evolved *S. aureus* populations regardless of treatment or lifestyle. **(A)** Average population size as determined by CFU/mL across five independently evolved replicates after 1 and 11 days of treatment exposure. **(B)** Average population size as determined by CFU/bead across all five independently evolved replicates after 1 and 11 days of treatment exposure. **(C)** Biofilm growth inhibition quantified using crystal violet (CV) staining. 10% v/v *B. subtilis* 6D1 CFE was applied to *S. aureus* populations at T0 and incubated statically at 37 °C for 24 hours. Residual biofilm biomass was quantified by CV staining and averaged by treatment/lifestyle. **(D)** Biofilm disruption quantified using CV staining. Either PBS or 10% v/v *B. subtilis* 6D1 CFE was applied to *S. aureus* 24-hour biofilms and incubated at 37°C shaking at 100 RPM for 2 hours. Residual biofilm biomass was quantified by CV staining and averaged by treatment/lifestyle. Differences were analyzed using a 2-tailed unpaired t-test where (****) P<0.0001, (***) P<0.001, (**) P<0.01 (*) P<0.05

To investigate how *S. aureus* adapted to the antibiofilm activity of *B. subtilis* 6D1 CFE, biofilm inhibition and disruption assays were performed to identify whether evolved lineages developed resistance to the antibiofilm activity of *B. subtilis* 6D1 CFE. When applied to *S. aureus* populations at time T0 to test biofilm prevention, CFE (10% v/v) reduced mean biofilm formation of all evolved lineages and the ancestor (Fig 2C). We also tested the effects of CFE on established biofilms to determine if CFE prompted biofilm disruption. Compared to biofilms treated only with PBS, CFE increased biofilm disruption against all *S. aureus* lineages (Fig 2D). Collectively, these results demonstrate *B. subtilis* 6D1 CFE maintains biofilm inhibition and disruption activity against all *S. aureus* lineages, regardless of long-term CFE exposure or lifestyle condition. These findings indicate that after ∼73 generations, *S. aureus* remained susceptible to the antibiofilm activity of CFE. However, in the absence of Bs CFE, significantly higher biofilm biomass was observed in the Bs CFE biofilm-evolved lineage compared to all other conditions (Fig. 2C). These data suggest Bs CFE biofilm-evolved lineages, despite exhibiting smaller population sizes when grown in the presence of CFE (Fig. 2B), may have adapted a slightly stronger biofilm phenotype in the absence of CFE.

### S. aureus populations evolved in the presence of B. subtilis 6D1 aggregate, but aggregation can still be inhibited by Bs CFE

Biofilms are typically defined by their ability to attach to surfaces(55), however there is a great deal of phenotypic variation that occurs throughout the biofilm growth cycle that can contribute to the development of these surface-attached microbial communities(56). Auto-aggregation is an additional biofilm phenotype where cells adhere to each other rather than a surface(43), and like mature biofilms, aggregate formation also improves *S. aureus* survival in the host and increases antibiotic tolerance(57, 58). Therefore, sedimentation assays were performed to identify if an increase in aggregate-type biofilms evolved in any of our *S. aureus* lineages. Control biofilm-evolved lineages exhibited faster and higher total aggregation compared to the ancestor and control planktonic-evolved lineages after 8 hours and 24 hours (Fig. 3A). Interestingly, both Bs CFE-evolved biofilm and planktonic lineages aggregated at similar rates (Fig. 3B). In fact, Bs CFE planktonic-evolved populations achieved significantly more aggregation after only four hours compared to control planktonic-evolved populations. No differences in percent auto aggregation were observed between control and Bs CFE-evolved biofilm lineages at any of the timepoints tested. Similar to our previous biofilm inhibition and disruption experiments (Fig. 2C-D), the presence of 10% v/v CFE also inhibited auto aggregation in all treatment lineages (Fig. 3C-D). Interestingly, Bs CFE biofilm-evolved lineages were significantly more susceptible to the effects of CFE at all time points compared to the ancestor (Fig. 3D). These data demonstrate that *S. aureus* evolved in a biofilm or in the presence of CFE with antibiofilm activity may actually adapt to increase aggregation, an often-overlooked stage of biofilm formation(59).

**Figure 3.**
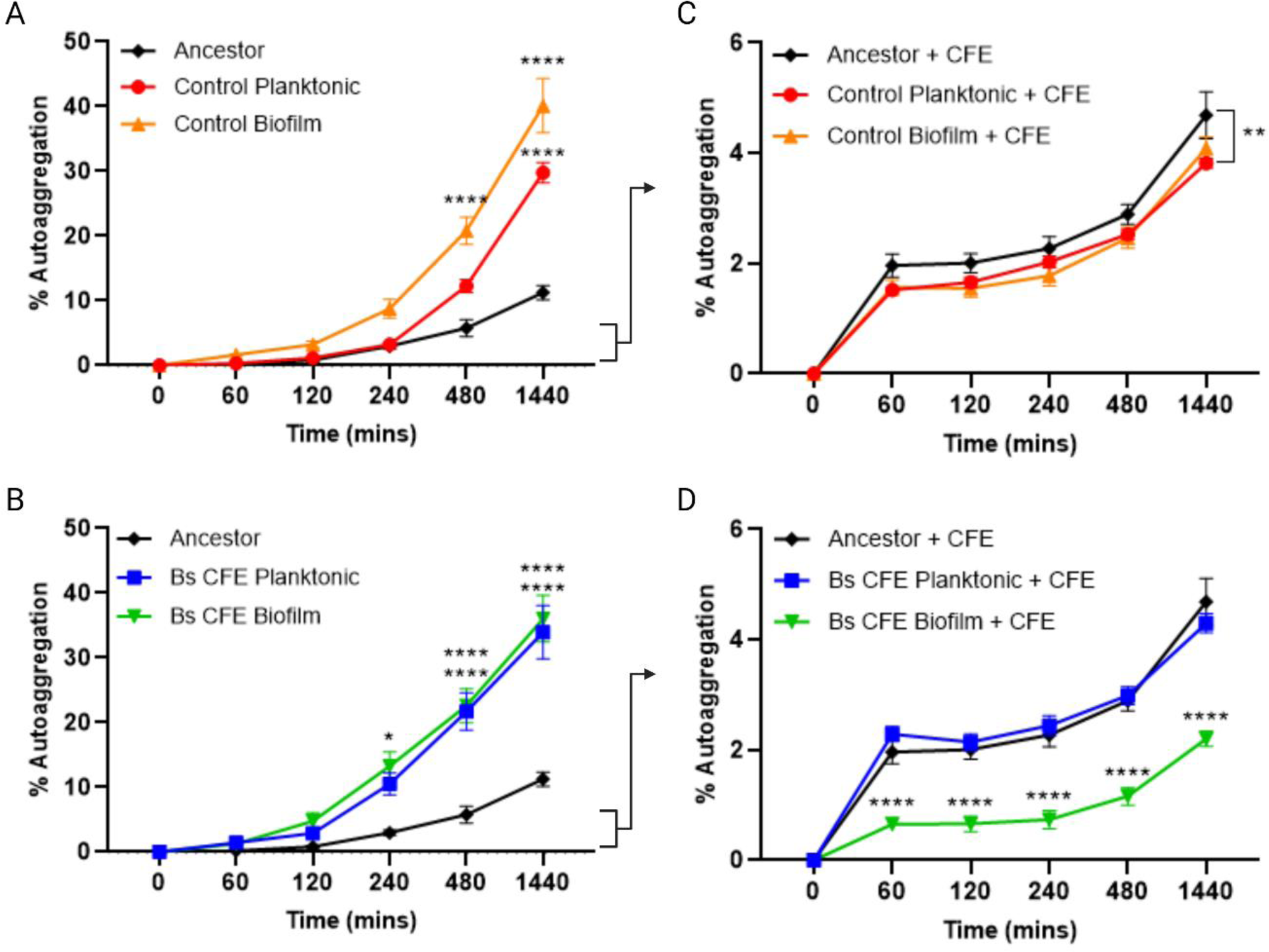
*B. subtilis 6D1 CFE* inhibits *S. aureus* auto-aggregation. Auto-aggregation rate of **(A)** control lineages and **(B)** Bs CFE exposed lineages averaged across replicates by treatment/lifestyle. Auto aggregation rate of **(C)** control lineages and **(D)** Bs CFE exposed lineages in the presence of 10% v/v *B. subtilis* 6D1 CFE averaged across replicates by treatment/lifestyle. Differences were analyzed using two-way ANOVA followed by Tukey’s multiple comparisons where (****) P<0.0001, (***) P<0.001, (**) P<0.01 (*) P<0.05 compared to the ancestor.

### S. aureus populations evolved in the presence of B. subtilis 6D1 CFE are less competitive in a biofilm environment against B. subtilis 6D1

To assess how these aggregative phenotypes affected competitive fitness, planktonic and biofilm competitions were performed between *S. aureus* lineages and *B. subtilis* 6D1. Planktonic competitions revealed control-evolved lineages were less competitive against *B. subtilis* 6D1 than the ancestor (Fig. 4A). Bs CFE planktonic-evolved lineages were more competitive than the planktonic-evolved untreated lineages. In this instance, the mean selection rate increased from −5.018 ± 0.84 for control planktonic-evolved lineages to −1.95 ± 0.51 for Bs CFE planktonic-evolved lineages. Mean selection rates also increased from −2.91 ± 0.78 to −1.24 ± 0.46 for control and Bs CFE treated biofilm-evolved lineages, respectively, however these differences were not statistically significant. These data indicate media adaptations caused a large competitive deficit in *S. aureus* lineages that manifest during competition against *B. subtilis* 6D1, but long-term exposure to CFE appears to slightly recover a portion of that fitness defect.

**Figure 4:**
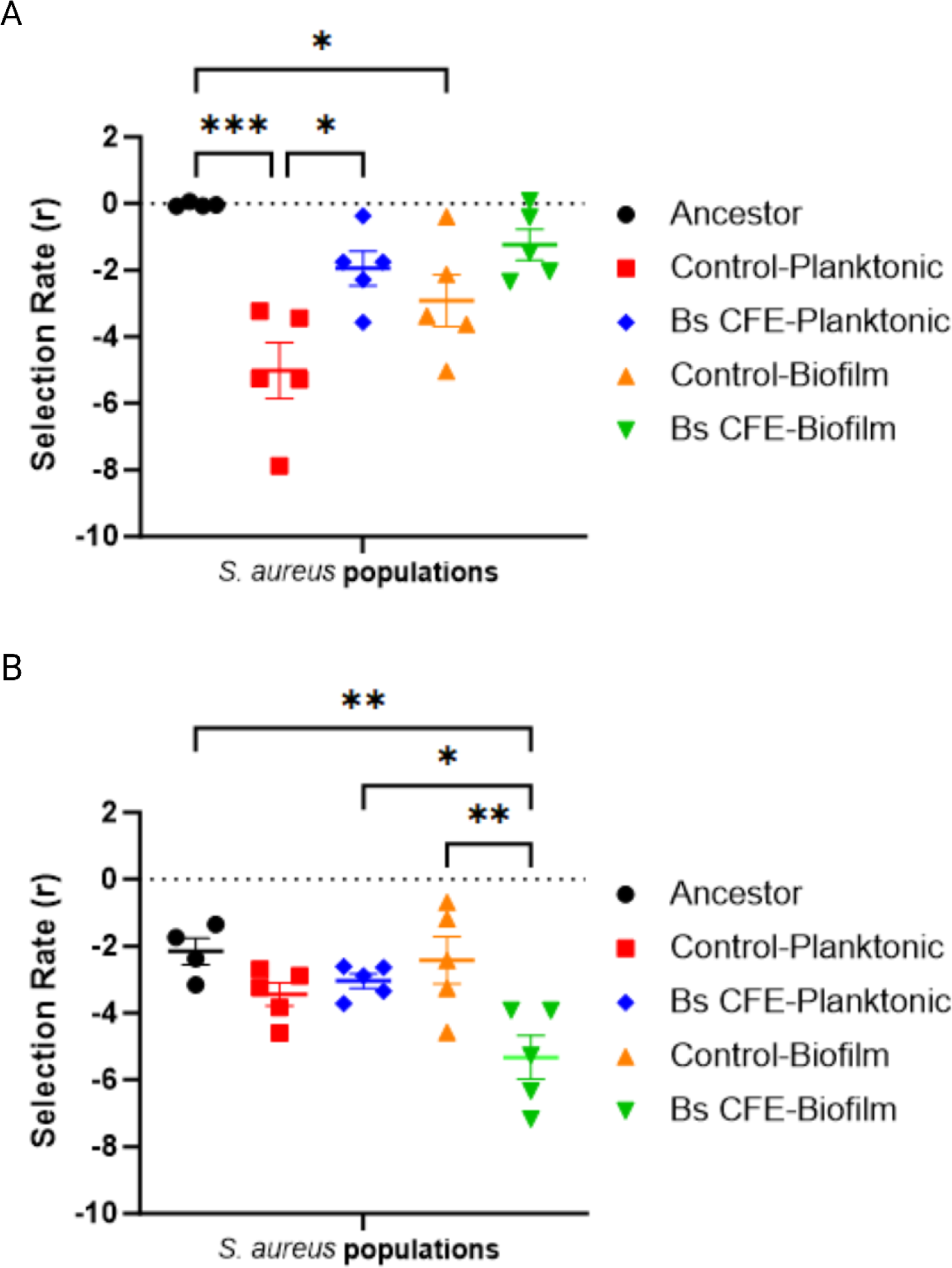
*S. aureus* populations evolved in the presence of *B. subtilis* 6D1 CFE are less competitive in a biofilm environment against *B. subtilis* 6D1. **(A)** Planktonic competition between *B. subtilis* 6D1 and independently evolved *S. aureus* populations. **(B)** Biofilm competition between *B. subtilis* 6D1 and independently evolved *S. aureus* populations. Each circle represents a single population tested in triplicate. Competitions with the ancestor were performed in quadruplicate using four separately prepared *S. aureus* ancestor cultures. Differences were analyzed using one-way ANOVA followed by Tukey’s multiple comparisons where (***) P<0.001; (**) P<0.01; (*) P<0.05

Unlike planktonic competitions, our biofilm competition data revealed only Bs CFE biofilm-evolved lineages were less competitive compared to the ancestor (Fig. 4B). These lineages (−5.32 ± 0.65) were also less competitive than both similarly evolved control biofilm-evolved lineages (−2.42 ± 0.70) and Bs CFE planktonic-evolved lineages (−2.64 ± 0.21). Therefore, in addition to being more susceptible to the anti-aggregation activity of CFE, our data shows Bs CFE exposed biofilm-evolved lineages are also less competitive against *B. subtilis* 6D1 in a biofilm. Together, these data strengthen the notion that CFE-mediated adaptations acquired by *S. aureus* lineages may be detrimental to their ability to proliferate or stably coexist in biofilm communities with *B. subtilis* 6D1 or its mixture of secondary metabolites.

### S. aureus populations evolved in the presence of B. subtilis 6D1 CFE are less virulent than the ancestor

To assess whether fitness defects observed in our competition experiments correlated with an altered virulence potential, the *S. aureus* ancestor and all evolved lineages were applied to Vero (CCL81) cells and evaluated for cytotoxic effects via lactate dehydrogenase (LDH) release(48). Vero cell toxicity for both Bs CFE biofilm and planktonic-evolved lineages was approximately 30% less cytotoxic than the ancestor (Fig. 5A). LDH release also trended lower in similarly evolved control populations compared to the ancestor, yielding 56.31 ± 6.44% and 55.22 ± 4.88% cytotoxicity for planktonic and biofilm lineages, respectively. Though no difference in LDH release was observed between evolutionary conditions, unhealthy Vero cell phenotypes appeared less frequently and were less pronounced in cell lines infected with Bs CFE-evolved lineages than control-evolved lineages (Fig. 5B). These data suggest that adaptation to the growth media, rather than long-term exposure to *B. subtilis* 6D1 CFE, may be a stronger indicator of reduced virulence potential of *S. aureus*.

**Figure 5:**
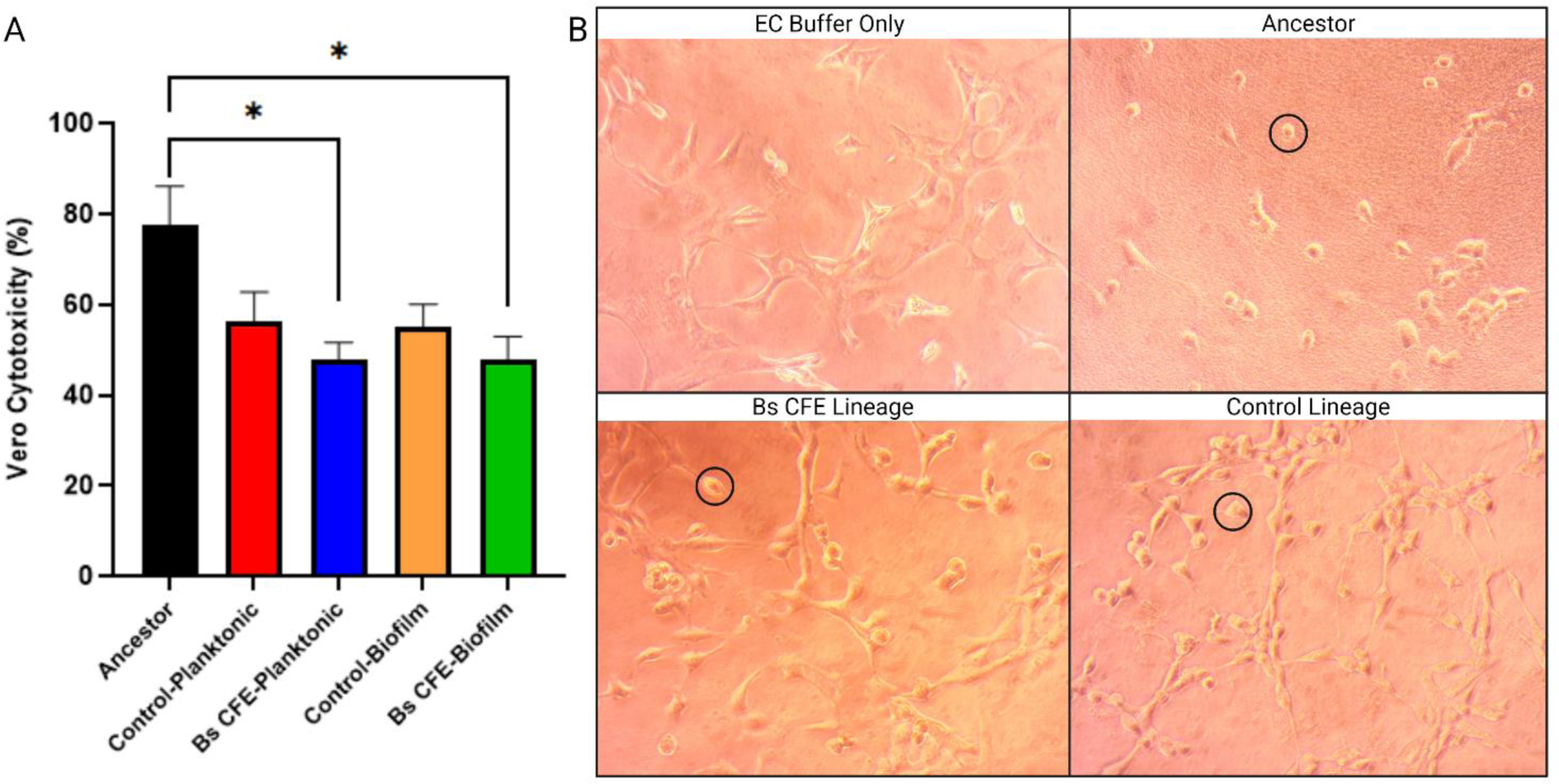
*S. aureus* populations evolved in the presence of *B. subtilis* 6D1 CFE are less virulent than the ancestor. **(A)** Vero cell cytotoxicity measured via lactate dehydrogenase release after 2-hour infection with *S. aureus* populations. **(B)** Vero cell morphology observed at 20X magnification after 2-hour exposure to representative *S. aureus* populations evolved in the presence or absence of Bs CFE. Black circles highlight blunted morphologies indicative of unhealthy Vero cells. Independently evolved populations were assayed in triplicate and averaged per treatment-lifestyle. Data were normalized to Vero cell lysis positive control. Differences were analyzed using one-way ANOVA followed by Tukey’s multiple comparisons where (*) P<0.05.

### Population sequencing reveals treatment and lifestyle-associated molecular targets of S. aureus evolution

Whole population genome sequencing was performed to identify genetic mutations that might explain the phenotypic characteristics observed across *S. aureus* evolved lineages. Collectively, in addition to discernible parallelism observed across treatment replicates, our results revealed mutations incurred by *S. aureus* ATCC 29213 after ∼73 generations followed treatment and lifestyle associated patterns (Fig. 6). Mutations identified in the chromosome affected genes associated with cell surface proteins – namely efflux pumps, competence proteins, clumping factors, and peptidoglycan synthesis enzymes. Mutations were also detected in genes related to RNA polymerase assembly and function. Most notably, we observed media-specific parallelism across all treatments, including controls, related to DNA polymerase function. All lineages incurred intergenic point mutations (A → T) at 100% frequency 174 bp downstream of *dnaN*, which encodes a DNA beta sliding clamp, and 207 bp upstream of *yaaA,* which encodes a S4 domain containing protein. In *E. coli*, YaaA functions as a DNA-binding protein that plays an important role in DNA maintenance during oxidative stress(60), and DNA beta clamps display a great deal of structural diversity to ensure consistant DNA polymerase function under different physiological conditions(61). Because reactive oxygen species can act as mutagens(62, 63), it is possible this mutation evolved to maintain DNA polymerase function in response to elevated reactive oxygen species produced when cells were grown in rich media(64, 65).

**Figure 6:**
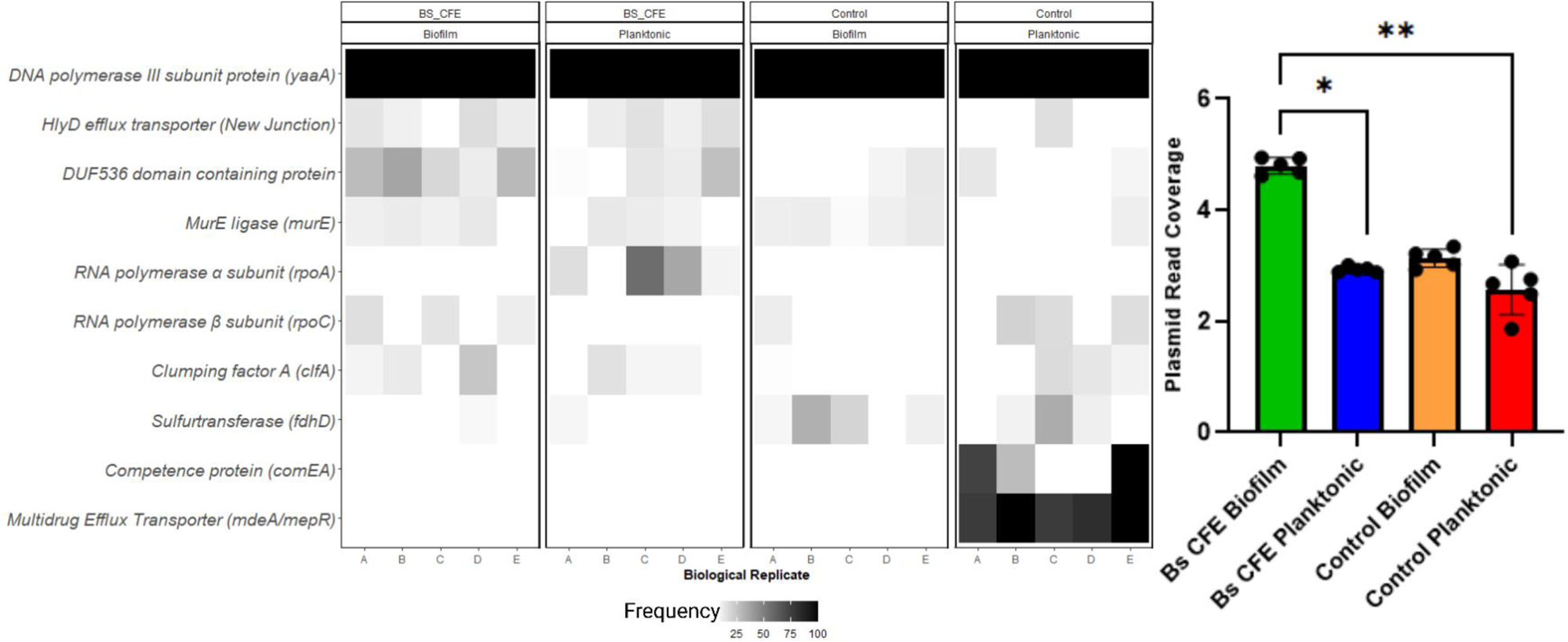
Population sequencing reveals treatment and lifestyle-associated molecular targets of *S. aureus* evolution. **(Left)** Mutations identified by whole-population genome sequencing of *S. aureus* lineages evolved in the presence and absence of 10% v/v *B. subtilis* 6D1 CFE. Five populations per treatment were sequenced after 11 days of experimental evolution. Mutations identified in the evolved populations that were not present in the ancestor are denoted by shaded black boxes. Shading indicates the frequency of mutation detection in the entire evolved population. Mutations depicted are a subset of all mutations identified; gene mutations were selected if they were observed in at least 3 lineages from any of the evolutionary treatment conditions. **(Right)** Mean plasmid read coverage of each *S. aureus* population (black dot) was normalized to the mean chromosome read coverage and averaged by treatment/lifestyle. Differences were analyzed using a non-parametric Kruskal-Wallis test followed by Dunn’s multiple comparisons where (**) P<0.01; (*) P<0.05

Interestingly, control planktonic-evolved lineages had the clearest lifestyle-dependent mutational parallelism. Mutations in competence and multi-drug efflux pump genes were observed in this group. All replicates had at least one high frequency (>76.2%) mutation in multidrug efflux pump regulation genes, *mdeA* or *mepR*. A nonsense mutation yielding a premature stop codon was found in *mepR*, at residue E72, which truncates MepR 67 amino acids shorter than the full-length protein. This truncation likely abolishes protein activity since hydrophobic residues L76, I77, L93, I98, L100 and V101 that make up the alpha side chain are critical for function(66). Though no mutations occurred within the *mdeA* coding region, single point mutations (C → A) were identified in four of five lineages 84 bp upstream of the *mdeA* translation start site (Table S3). We predicted these putative loss of function mutations in efflux pumps might increase antibiotic susceptibility in these lineages, and both KB-disc diffusion and broth microdilution assays corroborated these assumptions (Fig. S2, Table S2). Interestingly, *S. aureus* populations propagated in the presence of *B. subtilis* 6D1 CFE also did not exhibit increased phenotypic resistance against any of the antibiotics tested.

Mutations in *murE*, which encodes an enzyme critical for maintaining peptidoglycan integrity, were identified predominantly in biofilm and Bs CFE-evolved lineages. A missense *murE* mutation (C → A) at residue 464 converted isoleucine to asparagine in all afflicted lineages except in one of the control biofilm lineages where a point mutation (C → A) converted residue 458 from a glycine to an arginine (Table S3). Both positions are located in the active site region of the MurE protein structure (Fig. S3) and involved in L-lysine substrate interactions(67). Mur ligases help to remodel and recycle peptidoglycan in *S. aureus*, and this recycling is known to drive bacterial aggregation and biofilm formation(68). Therefore, it is possible mutations identified in *murE* are contributing to the increased aggregation phenotype observed in biofilm-evolved and Bs CFE-evolved lineages. In addition to mutations affecting cell surface proteins, pathways affecting RNA polymerase (RNAP) structure and function were also identified in a treatment/lifestyle pattern. Specifically, *rpoA* missense mutations at residues A269T/V and R268C/H were only identified in Bs CFE planktonic-evolved lineages (Fig. 6). In *E. coli*, *rpo* mutations and a loss of general stress resistance were found to be driven by competing stress responses influenced primarily by overexpression of key regulatory sigma factors(69). It is known that *B. subtilis* 6D1 CFE upregulates unrelated regulatory pathways in *S. aureus*, including the SigB general stress response(37). Therefore, we postulate mutations in *rpoA* observed in CFE planktonic-evolved lineages are likely driven by multiple compounds in the CFE, creating different stress responses in *S. aureus*.

We also identified mutations in the circular 4.69 Kb plasmid. Specifically, a nonsense mutation in the HlyD family efflux transporter periplasmic adaptor subunit was identified in 80% of CFE-evolved lineages (Fig. 6, Table S3). Missense mutations affecting the DUF536 domain containing protein were also identified at higher frequency in Bs CFE biofilm-evolved lineages. These mutations were identified at residues G255C and S256C/I – the last two amino acid residues of this protein’s C-terminal domain (Table S3, Fig. S3). It has been proposed a conserved C-terminal DUF536 domain is required for theta-replicating plasmids in multiple bacterial species(70–72). In line with these findings, we observed higher mean coverage depth in mapped plasmid reads in Bs CFE biofilm-evolved lineages compared to planktonic-evolved lineages (Fig. 6). Therefore, it is possible the missense mutations identified in DUF536 may be involved in maintaining or inheriting essential plasmid genes during evolution. Plasmids play an important role in bacterial evolution by transferring niche-adaptive functional genes, and an increased plasmid copy number has been shown to confer a fitness benefit to cells exposed to different environmental stressors(73, 74). The increased number of plasmid reads detected in Bs CFE biofilm-evolved lineages might explain the increased biofilm formation observed when these populations were grown without CFE (Fig. 2C). However, acquisition or maintenance of plasmids may incur high fitness costs for the host cell. Therefore, it is possible maintenance of these plasmid replicons, combined with adaptations imparted by our media conditions yielded negative epistatic interactions that increased sensitivity of these lineages to CFE and reduced their competitive fitness against *B. subtilis* 6D1.

## Discussion

Many studies have explored the mechanisms by which probiotic strains inhibit pathogenic biofilms through QSI(75–80), however it is often unknown how these proposed anti-virulence strategies influence the adaptation and evolution of these bacterial populations. This study investigated how long-term exposure to probiotic-derived compounds with antibiofilm activity affect pathogen fitness, virulence, and antibiotic sensitivity. After ∼73 generations, *S. aureus* remained susceptible to *B. subtilis* 6D1 CFE antibiofilm activity (Fig. 2C-D). Furthermore, the CFE inhibited auto aggregation against all evolved *S. aureus* lineages regardless of treatment or lifestyle (Fig. 3B, 3D) despite the evolution of an increased auto aggregation phenotype (Fig. 3A, 3C). *B. subtilis* 6D1 also maintained higher competitive fitness against all *S. aureus* evolved lineages in biofilm environments (Fig. 4B). *S. aureus* lineages propagated in the presence of *B. subtilis* 6D1 CFE were less cytotoxic than the ancestor in a Vero cell line model of infection (Fig. 5) and did not develop phenotypic resistance to a variety of antibiotics targeting cell wall integrity, protein translation, and nucleic acid synthesis when compared against the broadly susceptible *S. aureus* ATCC 29213 ancestor(81, 82) (Fig. S2, Table S2).

Bacterial auto aggregation or clumping, in this case, is a process whereby bacteria physically interact with each other and settle to the bottom in a static liquid suspension. Despite an enhanced aggregative phenotype, *B. subtilis* 6D1 CFE maintained both antibiofilm and anti-aggregative activity against all *S. aureus* lineages regardless of evolutionary condition. These data suggest application of a mixture of CFE compounds might help to slow the emergence of a more resistant phenotype by applying multiple selective pressures on *S. aureus* simultaneously(31). Additionally, because QSI involves the use of small molecular peptide inhibitors/initiators, resistance may require a specific combination of mutations that would need a larger mutation supply to generate the narrow range of combinations of resistance determinants(83–85). For example, while ∼73 generations are estimated to sample a single mutation in every *S. aureus* ATCC 29213 gene, significantly more generations, higher population sizes, or increased mutation rates may be required to observe a subsequent mutation that needs the first to appear. Similar to biofilms, aggregate formation also improves *S. aureus* survival in the host and increases antibiotic tolerance(57, 58). *S. aureus* aggregates also exhibit higher metabolic activity than biofilm cells and harbor a slightly increased mutation frequency compared to planktonic cells(57). Recent work demonstrated that aggregates landing on a surface eventually outcompete a biofilm population arising from single cells(59), suggesting aggregates may have a fitness advantage when it comes to accessing nutrients. Thus, auto aggregation is believed to provide bacteria with the benefits of biofilm while maintaining mobility. Increased aggregation and biofilm formation have also been proposed as an adaptive strategy to help cells reduce the toxic effects of reactive oxygen species (ROS)(64, 86). Therefore, it is possible Bs CFE-evolved and control biofilm-evolved lineages developed enhanced aggregative phenotypes in response to elevated ROS associated with fast growth in rich media. The media adaptation observed in our control lineages also caused a fitness deficit when competed against *B. subtilis* 6D1 in a planktonic environment (Fig. 4A). While we observed some recovery of that fitness deficit in Bs CFE-evolved lineages, these results suggest strong epistatic effects imparted by the media conditions may have prevented the emergence of a more obvious CFE-evolved phenotype(87, 88). This might also suggest fitness tradeoffs occur between fast growth and competitive fitness – if pathogens grow quickly in nutrient rich environments, they may be more susceptible to incoming or existing competitors.

In support of this hypothesis, control planktonic-evolved lineages, harboring high frequency mutations in genes encoding major facilitator superfamily pump proteins MdeA and MepR(89), displayed significant fitness defects when competed against *B. subtilis* 6D1 in a planktonic environment. Mutations in *mdeA* and *mepR* may have also evolved to help planktonic cells alleviate the toxic effects of accumulating ROS achieved in fast growing nutrient rich conditions(90, 91). In *Pseudomonas aeruginosa*, elevated levels of ROS increases activation of antibiotic efflux pumps(92). In *S. aureus*, AgrA also operates as a redox-responsive transcription factor that is inhibited in response to elevated intercellular ROS(93, 94), therefore, it is possible that CFE-exposed populations did not develop *mdeA/mepR* mutations due to constitutive *agrA* expression brought on by exposure to the CFE(37). MepR is a transcriptional regulator of multidrug efflux pumps that represses expression of *mepA*(66). However, nonsense mutations identified in *mepR* at residue E72 likely truncated this protein and abolished or reduced transcriptional repression of *mepA*. Additionally, mutations identified −84 bp upstream of *mdeA* might also contribute to increased *mdeA* transcription and subsequent pump activity since point mutations previously identified at this position prompted *mdeA* overexpression and increased resistance to a variety of MdeA substrates(95). While both *mdeA* and *mepR* mutations should have contributed to an increased antibiotic resistance phenotype, we did not observe broad antibiotic resistance in these lineages (Fig. S2, Table S2). One potential explanation may be due to the overall reduced fitness of planktonic-evolved lineages in the starch-based MH media used to assess antibiotic sensitivity (Fig. S1). MepR activity is induced by multiple antimicrobial peptides(66), and the potency of these peptides is increased in planktonic TSB cultures compared to MHB cultures(96). Therefore, the use of MH media may have been inadequate to detect differences in MepA induced phenotypic resistance in these lineages. Despite this, *mdeA/mepR* mutations detected in these lineages may have contributed to the reduced fitness observed in our planktonic competition experiments (Fig. 4A).

Previous work in our lab established *B. subtilis* 6D1 CFE upregulates *agr, RNAIII*, and *saeR* in *S. aureus*(37), and overexpression of these regulators also increases transcription of peptidoglycan hydrolases *lytM, ssaA, sceD*(97). Peptidoglycan hydrolases and Mur ligases collectively help to remodel and recycle peptidoglycan in *S. aureus*, and peptidoglycan recycling is known to drive bacterial aggregation and biofilm formation(68). In our evolution experiment, mutations affecting MurE occurred primarily at I464 (Table S3) and predominantly in Bs CFE and control biofilm-evolved lineages (Fig. 6). Previous work demonstrated that point mutations affecting the MurE I464 residue impairs structural stability in the active site(67), and leads to increased peptidoglycan recycling and turnover(98–100). Based on these data, it is possible that mutations in *murE* I464 identified in our study contributed to peptidoglycan modifications that increased aggregation. A number of small molecule inhibitors designed to inhibit aggregation have also shown an ability to reduce *S. aureus* virulence(101–103), and we have previously demonstrated *B. subtilis* 6D1 CFE reduces *S. aureus* toxicity in a concentration dependent manner(37). Therefore, we investigated whether CFE-evolved lineages had adapted to overcome this anti-virulence activity by developing a more cytotoxic phenotype. Interestingly, CFE-evolved lineages were less cytotoxic compared to the ancestor (Fig. 5), suggesting long-term exposure to CFE did not increase *S. aureus*-induced toxicity in our infection model. However, since cytotoxicity also trended lower in control-evolved lineages, it is likely this reduced virulence can be attributed to adaptations to the growth media.

Expression and secretion of *S. aureus* virulence factors are tightly controlled by multiple regulatory systems, however almost all rely on functional RNA polymerases(104). RNA polymerase (RNAP) is the key enzyme responsible for transcription of DNA into RNA, and natural products produced by other bacteria have already demonstrated that it is possible to inhibit RNA polymerase function in *S. aureus*(105). In our study, missense mutations in *rpoA* were found only in Bs CFE planktonic-evolved lineages while *rpoC* missense mutations were identified in Bs CFE biofilm-evolved and control planktonic-evolved populations (Fig. 6). Bacterial RNAP core enzyme consists of several subunits: the α (*rpoA*), β (*rpoB*), β′ (*rpoC*), and the ω (*rpoZ*) subunit. As a form of adaptive evolution, the genes encoding RNAP subunits acquire mutations in response to virtually any selection pressure including media composition, laboratory domestication, antibiotic exposure, and oxidative stress(106–111). The α subunit encoded by *rpoA* plays an important role in RNAP assembly since its dimerization is the first step in the sequential assembly of subunits to form the holoenzyme(112). This core enzyme is conserved across the bacterial kingdom, however members of the Firmicutes contain an additional δ (*rpoE*) subunit(113). The δ subunit is essential for cell survival when facing a competing strain in a changing environment, and although δ is not essential, it is vital for the cell’s ability to rapidly adapt and survive in nature(114). In fact, *S. aureus* mutants where δ or ω fail to bind to the β’ subunit have diminished toxin activity and abundance, are less virulent in a murine model of infection(115), and harbor an impaired ability to resist stress and maintain biofilm structure(116). Therefore, *rpo* missense mutations identified in our experiment provide additional evidence that help explain the reduced virulence and competitive fitness of evolved lineages.

*B. subtilis* 6D1 CFE does not kill *S. aureus*(37), and since quorum sensing interference is not believed to induce strong selection pressure, it is possible the increased antibiotic sensitivity and decreased virulence observed in our Bs CFE-evolved populations can be attributed to the fitness cost of harboring excess plasmid replicons(73) (Fig. 6). Mutations in plasmid-associated genes were also found more frequently and in higher abundance in CFE-exposed lineages (Fig. 6). Conversely, lower plasmid read coverage observed in control populations may also help to explain the competitive deficiencies observed during planktonic competition against *B. subtilis* 6D1 (Fig. 4A). Taken together, these data suggest an increase in plasmid-borne traits may be responsible for improved planktonic fitness observed in CFE-evolved lineages. Therefore, it is possible this plasmid plays an important role in mediating fitness and virulence in stressful environments. In fact, *racA*, the gene reported to transcribe DUF536, is significantly correlated with gut-associated *S. epidermidis* isolates(117), and loss of the plasmid harboring *racA* rendered a clinical MRSA strain Agr defective, but hypervirulent(118). In line with these findings, *S. aureus* QS signal-blind mutants possess a significant survival advantage *in vivo*(119), and a separate study reported that *S. aureus* strains that caused the most severe disease were the least toxic due, in part, to within-host fitness differences(120). Despite previous clinical trials showing a reduction in *S. aureus* colonization after supplementation with a *B. subtilis* probiotic(34), future studies should aim to investigate how residual *S. aureus* populations evolve in these host-associated systems. Still, our study provides the first evidence that long-term exposure to a collection of probiotic derived compounds harboring QSI activity, like those found in *B. subtilis* 6D1 CFE, can inhibit multiple *S. aureus* virulence mechanisms without increasing resistance to antibiotics.

## Author Contributions

K.R.H and C.W.M. conceptualized and supervised the study. K.R.L. contributed to the study design. K.R.L., E.S, and J.K. conducted the investigation and performed laboratory analysis. K.R.L. performed the formal data analysis and its visualization and wrote the original draft. All authors edited and approved the final manuscript.

## Acknowledgements

We would like to thank Grace Schmaling and Katie Zingelman for their laboratory assistance and Seqcenter for sequencing technical support. We would also like to acknowledge Microbial Discovery Group for their continued support of student research. This work was funded, in part, by the Department of Defense Office of Army Research Grants W9132T-22-2-0001 and W9132T-23-2-0003. Any opinions, findings, and conclusions or recommendations expressed in this material are those of the authors and do not reflect the official policy or position of the Department of the Army, Department of Defense, or the U.S. Government.

## Supplementary Tables & Figures

**Table S1.** Evolved lineages exhibit improved growth rates in evolution media compared to the ancestor.

**Supplementary Tables & Figures:**
|  |  |  |  |
| --- | --- | --- | --- |
| **Treatment/Lifestyle** | **Slope (R<sup>2</sup>)** | **Mean Exponential Growth Rate ($\mu$)** | **Mean Doubling Time (mins)** |
| Ancestor | 0.9912 | 0.64 | 84.28 |
| Control Planktonic | 0.9938 | 0.93 | 63.42 |
| Bs CFE Planktonic | 0.9967 | 0.95 | 62.2 |
| Control Biofilm | 0.9945 | 0.98 | 60.46 |
| Bs CFE Biofilm | 0.9963 | 0.89 | 65.46 |

**Table S2.** Minimum inhibitory concentrations of evolved S. aureus lineages exposed to various antibiotics.

| Population | Antibiotic MIC (µg/mL) |  |  |  |  |  |  |  |
| --- | --- | --- | --- | --- | --- | --- | --- | --- |
|  | Gentamicin | Tetracycline | Rifampin | Erythromycin | Sulfa-Trim | Cefotaxime | Ceftazidime | Vancomycin |
| Ancestor | 2.5 | 15 | 5 | 30 | 23.75x1.25 | 1.87 | 30 | <1.87 |
| Control Planktonic A | 2.5 | 7.5 | 5 | 30 | 23.75x1.25 | 3.75 | 15 | <1.87 |
| Control Planktonic B | 1.25 | 7.5 | 5 | 15 | 23.75x1.25 | 3.75 | 15 | <1.87 |
| Control Planktonic C | 2.5 | 7.5 | 2.5 | 30 | 23.75x1.25 | 3.75 | 30 | <1.87 |
| Control Planktonic D | 1.25 | 7.5 | 5 | 30 | 23.75x1.25 | 3.75 | 30 | <1.87 |
| Control Planktonic E | 1.25 | 15 | 5 | 30 | 23.75x1.25 | 1.87 | 30 | <1.87 |
| Bs CFE Planktonic A | 1.25 | 7.5 | 5 | 15 | 23.75x1.25 | 1.87 | 30 | <1.87 |
| Bs CFE Planktonic B | 1.25 | 7.5 | 2.5 | 15 | 23.75x1.25 | 0.93 | 30 | <1.87 |
| Bs CFE Planktonic C | 1.25 | 15 | 2.5 | 15 | 23.75x1.25 | 0.46 | 30 | <1.87 |
| Bs CFE Planktonic D | 1.25 | 15 | 2.5 | 15 | 23.75x1.25 | 1.87 | 30 | <1.87 |
| Bs CFE Planktonic E | 2.5 | 15 | 5 | 15 | 23.75x1.25 | 0.93 | 30 | <1.87 |
| Control Biofilm A | 2.5 | 7.5 | 5 | 30 | 23.75x1.25 | 1.87 | 15 | <1.87 |
| Control Biofilm B | 1.25 | 7.5 | 2.5 | 30 | 23.75x1.25 | 1.87 | 15 | <1.87 |
| Control Biofilm C | 1.25 | 7.5 | 2.5 | 30 | 23.75x1.25 | 3.75 | 15 | <1.87 |
| Control Biofilm D | 2.5 | 7.5 | 2.5 | 30 | 23.75x1.25 | 1.87 | 15 | <1.87 |
| Control Biofilm E | 1.25 | 7.5 | 5 | 30 | 23.75x1.25 | 3.75 | 15 | <1.87 |
| Bs CFE Biofilm A | 2.5 | 7.5 | 2.5 | 15 | 23.75x1.25 | 1.87 | 7.5 | <1.87 |
| Bs CFE Biofilm B | 1.25 | 7.5 | 2.5 | 15 | 23.75x1.25 | 0.93 | 7.5 | <1.87 |
| Bs CFE Biofilm C | 2.5 | 15 | 5 | 15 | 23.75x1.25 | 1.87 | 15 | <1.87 |
| Bs CFE Biofilm D | 2.5 | 7.5 | 5 | 15 | 23.75x1.25 | 1.87 | 15 | <1.87 |
| Bs CFE Biofilm E | 1.25 | 7.5 | 5 | 15 | 23.75x1.25 | 0.93 | 15 | <1.87 |

**Figure S1.**
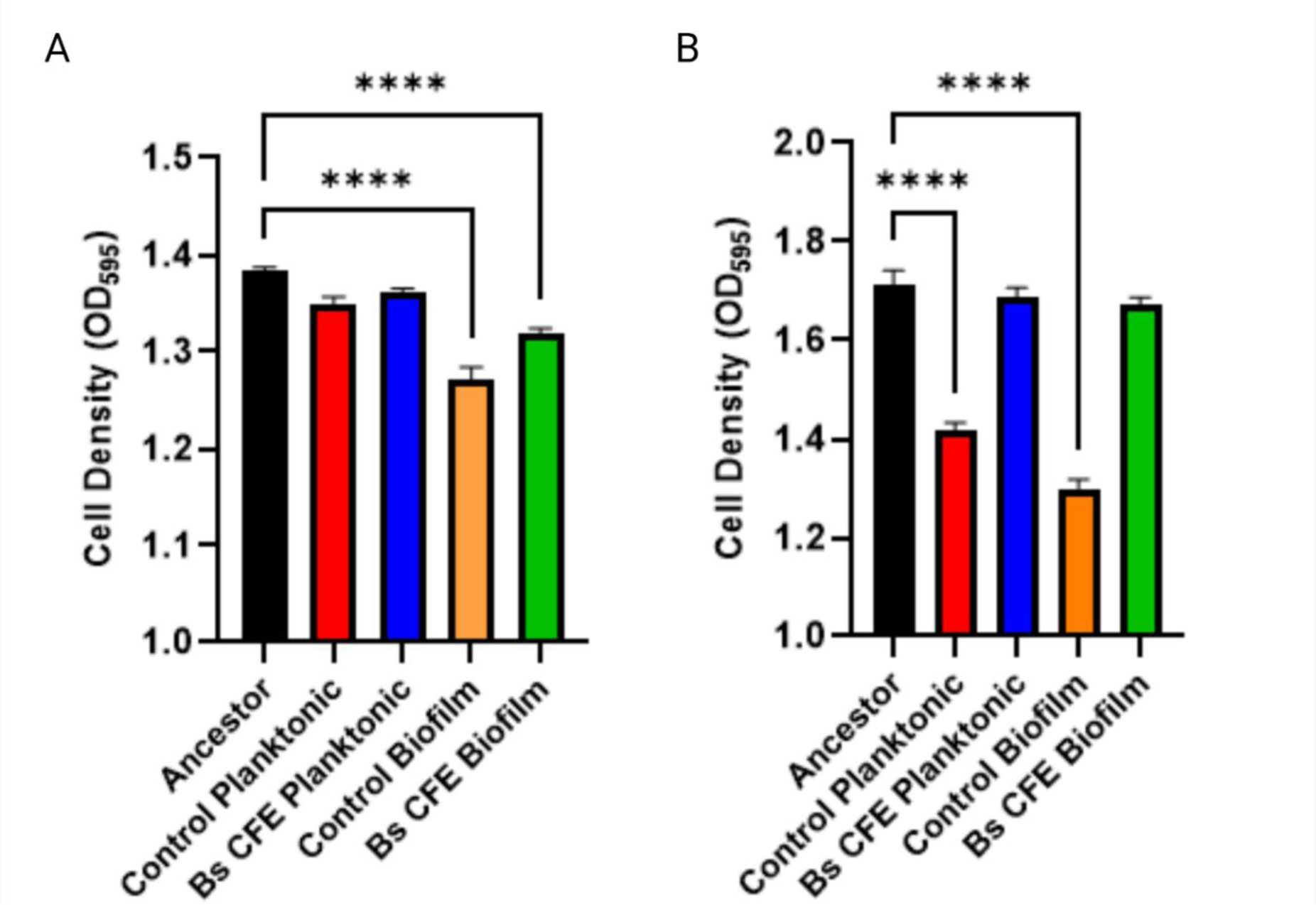
Media composition reveals treatment and lifestyle dependent 24-hour growth patterns. Planktonic cell density achieved after 24-hour incubation at 37°C in 1.5% TSB + 0.3% glucose **(A)** and MHB **(B)** averaged across replicates by treatment/lifestyle. (****) P<0.0001

**Figure S2:**
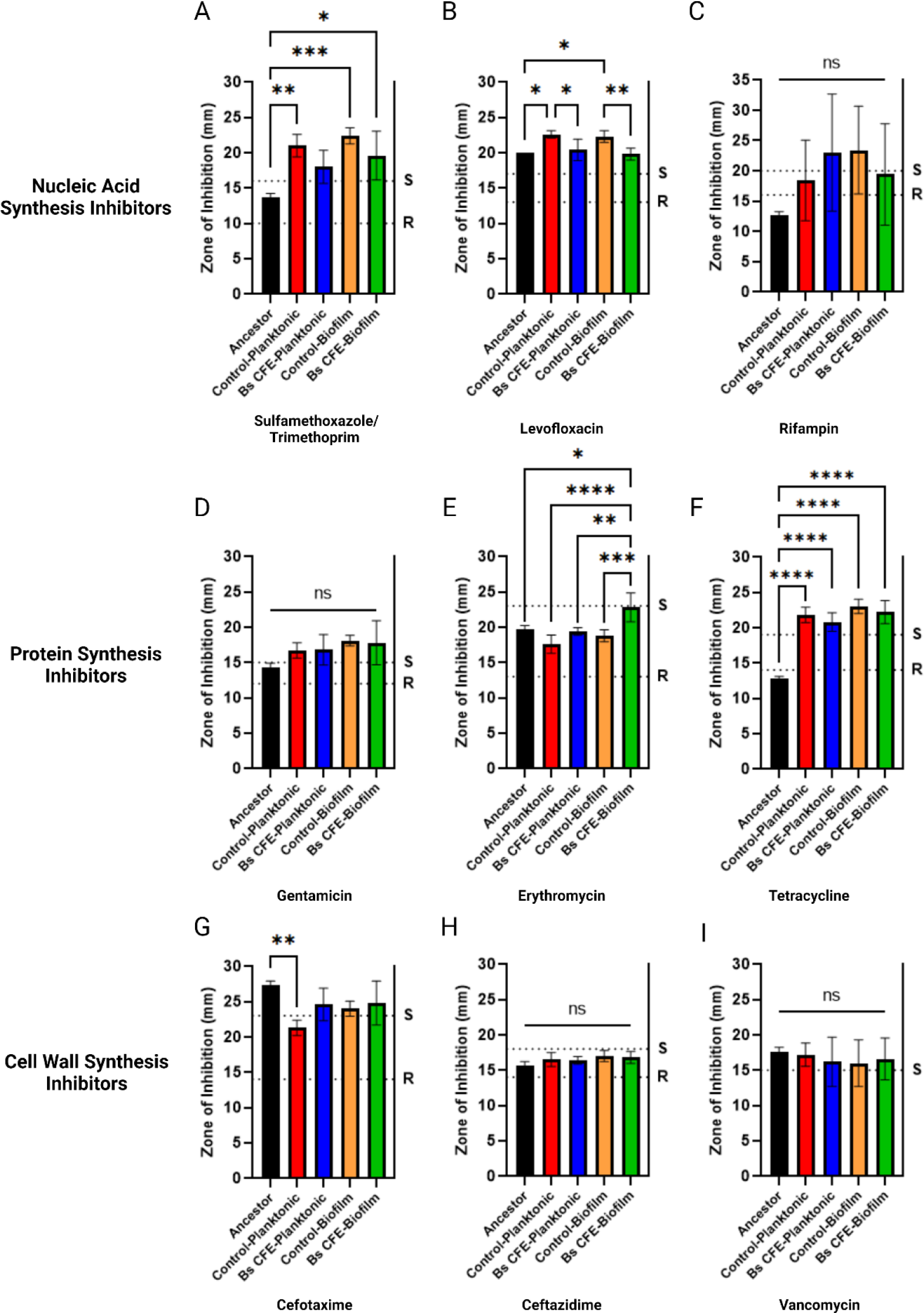
Antibiotic susceptibility testing of *S. aureus* lineages reveals treatment and lifestyle-associated phenotypic sensitivity patterns. KB disc diffusion assays were used to compare antibiotic sensitivities of the *S. aureus* 29213 ancestor and evolved populations. (S)usceptible and (R)esistance thresholds defined by Sensi-disc™ protocols for *Staphylococcus* are plotted as dashed lines. Bars represent mean ± SD averaged across replicates by treatment/lifestyle. Differences were analyzed using one-way ANOVA followed by Tukey’s multiple comparisons where (****) P<0.0001 (***) P<0.001; (**) P<0.01; (*) P<0.05

**Figure S3.**
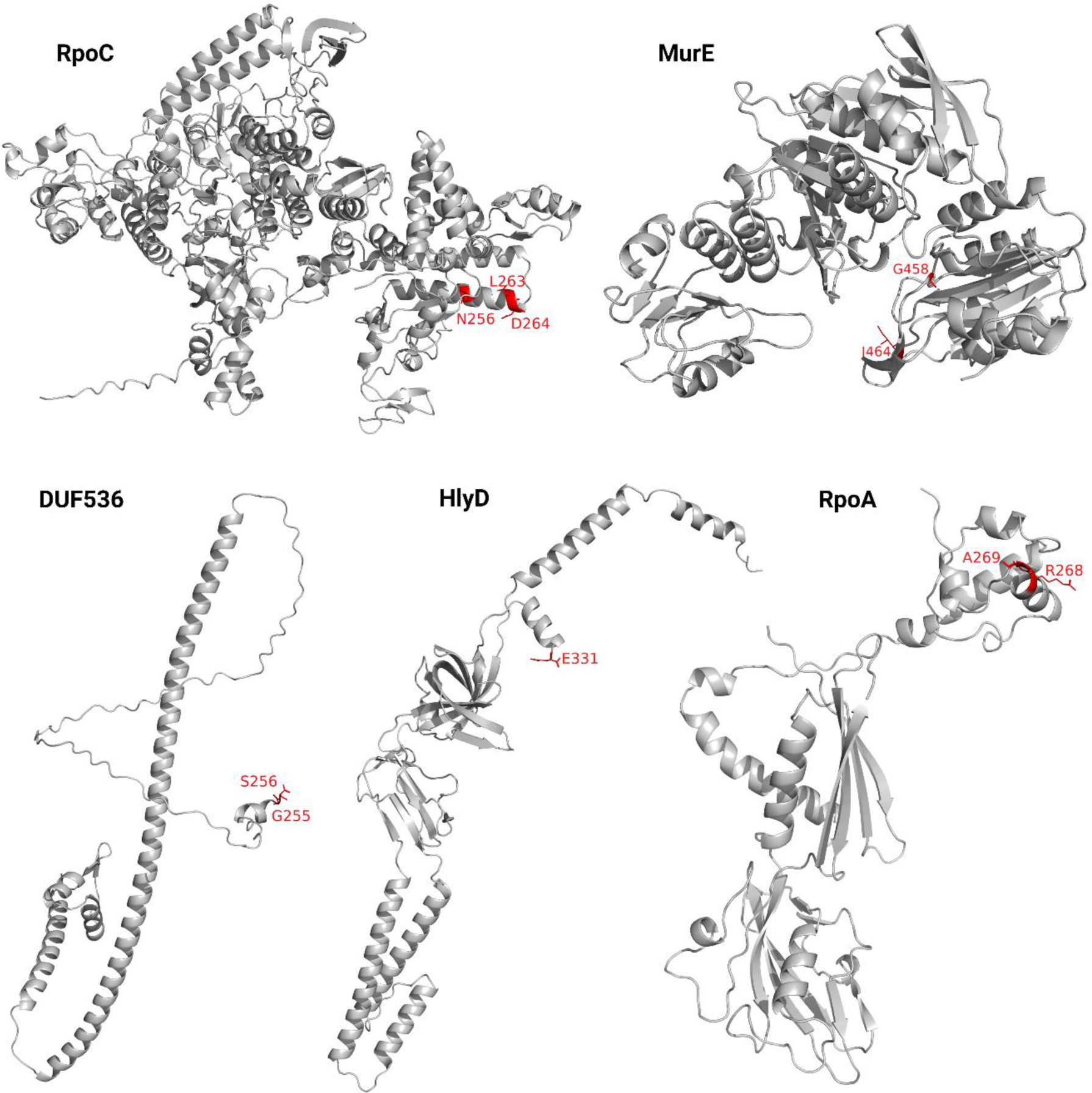
Missense mutations acquired during *S. aureus* ATCC 29213 evolution. Amino acid residues harboring missense mutations (red) were mapped onto AlphaFold predicted protein structures and visualized using Pymol.

**Figure S4.**
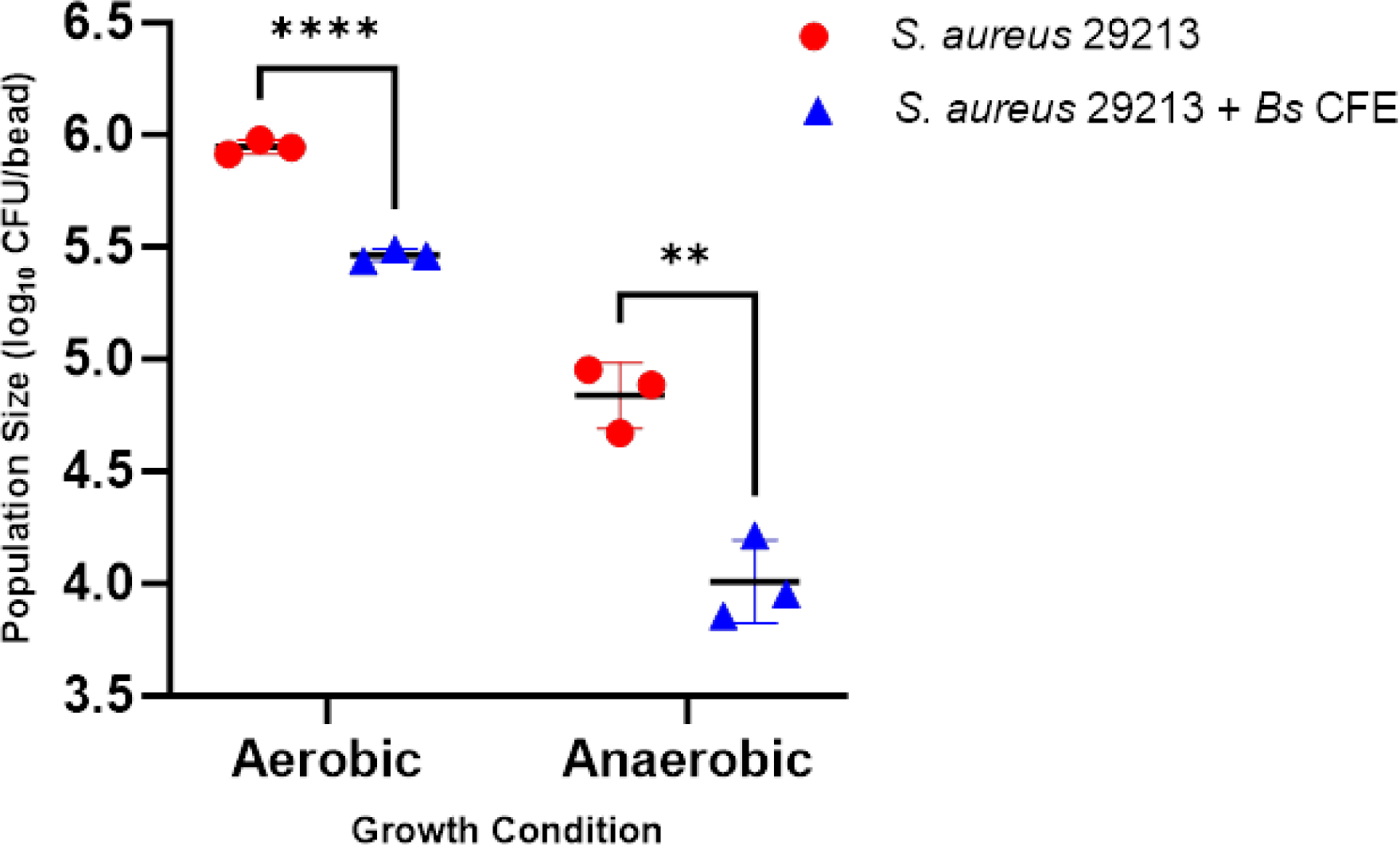
*B. subtilis* 6D1 CFE inhibits *S. aureus* biofilm growth on a polystyrene bead under aerobic and anaerobic conditions. *S. aureus* 29213 was inoculated into 5mL 1.5% TSB + 0.3% glucose in a glass test tube harboring a single polystyrene bead. These cultures were grown at 37°C for 24 hours in the presence (blue triangles) or absence (red circles) of 10% v/v *B. subtilis* 6D1 CFE (*Bs* CFE) under either aerobic or anaerobic conditions. Beads were removed from the culture media, gently rinsed in PBS, placed into 1mL sterile PBS, and sonicated at 60Hz for 10s prior to CFU enumeration. Biofilm growth inhibition was quantified across three independent experiments. Bars represent mean ± SD. Differences were analyzed using a 2-tailed unpaired t-test where (****) P<0.0001, (**) P<0.01

